# Neural correlates of perceptual biases in duration perception

**DOI:** 10.1101/2025.09.13.675931

**Authors:** Zahra Shirzhiyan, Stefan Glasauer

## Abstract

How we perceive a current event depends not only on its immediate context, but also on how our internal expectations are shaped by prior experience. In time perception, these expectations manifest as systematic biases, namely sequential dependence, where the current percept is influenced by the previous stimulus, and central tendency, the overestimation of short durations and underestimation of long ones. Both perceptual biases, corresponding to individual beliefs about stimulus generation, can vary substantially between participants. However, the neural correlates of these individual beliefs and their effects are unknown. Here, we investigate how these biases and their individual variations are reflected in neural responses in a duration reproduction task. Our EEG results show that in the frontocentral region, the Contingent Negative Variation (CNV) while experiencing the current stimulus depends on the previous stimulus regardless of whether sequential dependence is high or low. In contrast, in the right parietal region, CNV significantly correlated with the amount of sequential dependence. Central tendency was associated with frontocentral CNV amplitude and post-stimulus P2 components. A Bayesian model of time perception reproduced the observed neural dynamics, suggesting that internal estimates and expectations of stimulus offset are reflected in EEG responses. Our results demonstrate that both forms of perceptual bias, sequential dependence and central tendency, are reflected in neural activity while experiencing the ongoing stimulus, suggesting that both biases directly affect the measurement of time.

**New and Noteworthy:** Individuals differ in how they perceive magnitudes, exhibiting systematic biases, sequential dependence and central tendency, which reflect internal expectations about stimulus generation. Using EEG, we show that both behavioral biases are reflected in neural dynamics during stimulus encoding, which can be reproduced by a Bayesian model. The correspondence between perceptual biases and patterns of brain activity indicates that perceptual history and expectations directly influence how the brain measures and represents time.

## Introduction

The perception of current events is influenced by previous experiences in the immediate past. This phenomenon, which is referred to as sequential or serial dependence (Cicchini et al., 2024), has been found in visual perception (Fischer and Whitney, 2014; Pascucci et al., 2023), and also for perception of magnitudes such as loudness (Holland and Lockhead, 1968; Cross, 1973; Akrami et al., 2018), distance (Petzschner et al., 2015; Glasauer and Shi, 2022), or duration (Dyjas et al., 2012; Glasauer and Shi, 2022). Similarly, central tendency, the bias toward the mean duration in estimation tasks, has been widely demonstrated across various perceptual magnitudes and studied using both behavioral paradigms (Jazayeri & Shadlen, 2010; Shi et al., 2013; Glasauer & Shi, 2021) and neuroimaging approaches such as fMRI (Murai & Yotsumoto, 2016).

To explain sequential effects, various models have been proposed (Jesteadt et al., 1977; Petzschner and Glasauer, 2011; Dyjas et al., 2012) which share that the current perceptual estimate depends on the previous one. Using probabilistic Bayesian modeling we have shown that sequential dependence optimizes perceptual estimates when stimuli are auto-correlated over time, as for example in a random walk (Glasauer and Shi, 2021, 2022). Consequently, if participants believe that subsequent stimuli depend on each other, then their percepts will display sequential dependence even if stimuli are generated independently (Glasauer and Shi, 2022). Individual differences in perception (Glasauer, 2024) reflected in the strength of perceptual biases can thus be explained in part by differences in beliefs about how stimuli are generated in the world.

Here we aim to identify neural correlates of perceptual biases such as the sequential dependence to elucidate the mechanisms underlying perception. EEG studies have used duration perception paradigms to investigate neural signatures such as the Contingent Negative Variation (CNV) and event-related potentials (P2, P3, LPCt) (Ng and Penney, 2014; Wiener and Thompson, 2015; Kononowicz and Penney, 2016; Kononowicz et al., 2018). The on frontocentral electrodes CNV is a slow negative potential that develops after stimulus onset predominantly over frontocentral electrodes. It has been proposed that it could provide information regarding temporal memory reflecting readiness and expectancy (Walter et al., 1964; Pfeuty et al., 2005), or be related to higher order cognitive tasks like decision-making and motor preparation (Kononowicz and Penney, 2016; Kononowicz et al., 2018). The frontal CNV recently has been shown to be modulated by the previous stimulus, which supposedly results from dynamical expectations (Damsma et al., 2021).

Our current study investigates how individual differences in time perception (Glasauer and Shi, 2022) are reflected in neural correlates measured by EEG in a duration reproduction task (see Fig. 1A). We previously showed (Glasauer and Shi, 2021, 2022) that sequential dependence and central tendency, a bias leading to overestimation of small magnitudes and underestimation of large ones, can differ substantially between participants. Predictions of our Bayesian model (Glasauer and Shi 2022) for possible combinations of biases (Fig. 1B) are shown in Fig. 1C-E. The reproduced duration can show strong (1C) or weak sequential (1D) dependence independently of central tendency, except if central tendency is low (1E), in which case sequential dependence is also low. The model posits that central tendency is caused by reliance on prior information. This prior information can either be a summary statistic of the distribution of possible stimuli, or just the previous stimulus, or combinations of both. If the previous stimulus is used as prior information, i.e., as predictor for the current stimulus, then sequential dependence becomes high. If only the summary statistic is used, sequential dependence will be zero, but central tendency can still be high. If participants rely completely on sensory input and do not use prior information, then both central tendency and sequential dependence are close to zero. From the model simulations (see also Methods), we propose internal representations of model variables that possibly are reflected in neural activity: the estimate of the experienced duration during stimulus encoding in the production phase (Fig. 1C3-E3), and the instantaneous expectancy for stimulus offset (end of production phase) as derived from the prior probability density (Fig 1C4-E4). For example, if sequential dependence is strong (1C), the running estimate of time (1C3) would vary depending on the previous stimulus.

**Figure 1.**
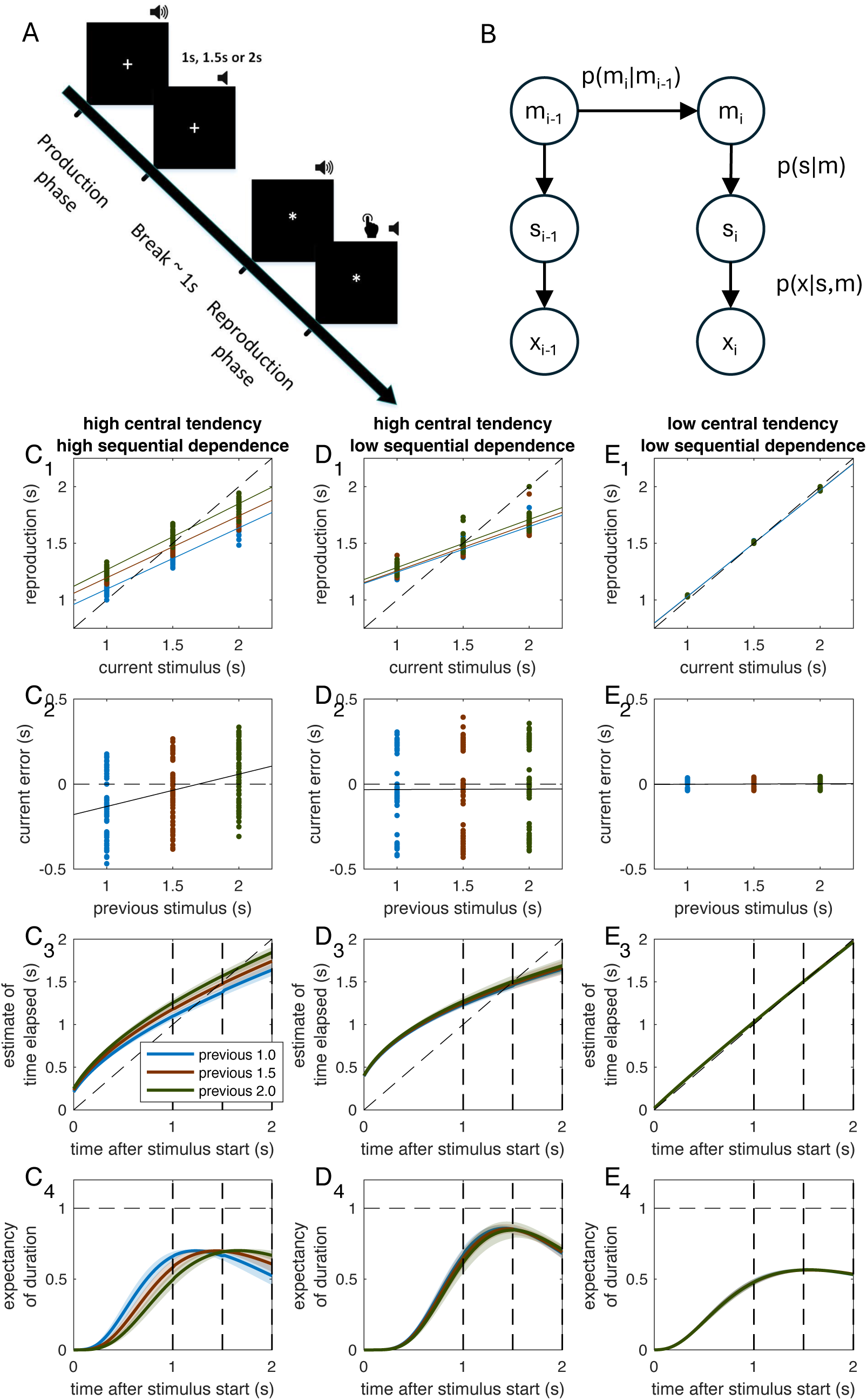
**A**: trial structure of the duration reproduction task employed in the current study (180 trials per subject, durations 1.0, 1.5, and 2.0 s). **B:** graphical depiction of the generative model of stimulus production (Glasauer and Shi, 2022). m_i_: mean of stimulus distribution at trial i, s_i_: stimulus duration at trial i, x_i_: noisy sensory measurement of stimulus duration. The current stimulus s_i_ is taken from a probability distribution whose mean m_i_ depends on the previous mean m_i-1_ with transition probability p(m_i_|m_i-1_). **C – E**: Model predictions for different combinations of perceptual biases. All durations in s. Colors denote the immediate previous stimulus. **C**: simulated participant with high central tendency and high sequential dependence. **D**: high central tendency but low sequential dependence. **E**: low central tendency and low sequential dependence. **C_1_, D_1_, E_1_**: current reproduction plotted over current stimulus duration, regression slope corresponds to 1 − central tendency. Each dot corresponds to one trial. **C_2_, D_2_, E_2_**: current error plotted over previous stimulus duration, regression slope corresponds to sequential dependence. **C_3_, D_3_, E_3_**: Model prediction for hypothesized running estimate of the elapsed duration from the start to the end of the production phase, plotted separately by previous stimulus. The curvature is caused by the model operating in the logarithmic domain. **C_4_, D_4_, E_4_**: Model prediction for hypothesized expectancy of stimulus duration. Expectancy is given as instantaneous prior probability density of predicted stimulus duration. For high sequential dependence (C), both the estimate of elapsed duration (C_3_) and the expectancy of stimulus end (C_4_) depend on previous stimulus duration.

To identify neural patterns reflecting individual differences, we grouped participants according to the strength of their individual biases. For example, we expected that if the frontal CNV reflects the internal estimate of elapsed duration, then the CNV’s dependence on previous stimulus duration (Damsma et al., 2021) would differ between participants with large vs. small sequential dependence, similar to the model prediction in Fig. 1C3 vs. D3. In addition to frontal activity, we investigated responses in the right parietal lobe, which has also been implicated in perception of time intervals and the encoding of temporal information (Bueti and Walsh, 2009; Merchant et al., 2013; Contò et al., 2025), but also in attention and decision making (Heilman and Abell, 1980; Agosta et al., 2017).

## Materials and Methods

### Participants

We performed a power analysis using G*Power 3.1 (Faul et al. 2009) to determine sample size using data from a previous study that evaluated the effect of the duration of a previously experienced time interval on the CNV during production of a current interval (Damsma et al. 2021; Fig. 2F, 22 participants). From the published figure we estimated that a difference in prior duration of 0.5 s would result in a CNV amplitude difference of about 0.8 – 1.0 μV with a standard error of the mean of about 0.5 μV. Given repeated measures with a correlation of 0.75 and a desired power of 0.95, the effect size was determined as 0.492 – 0.615 resulting in a required sample size of 31 – 47. To be on the safe side, we opted for 48 participants.

**Figure 2.**
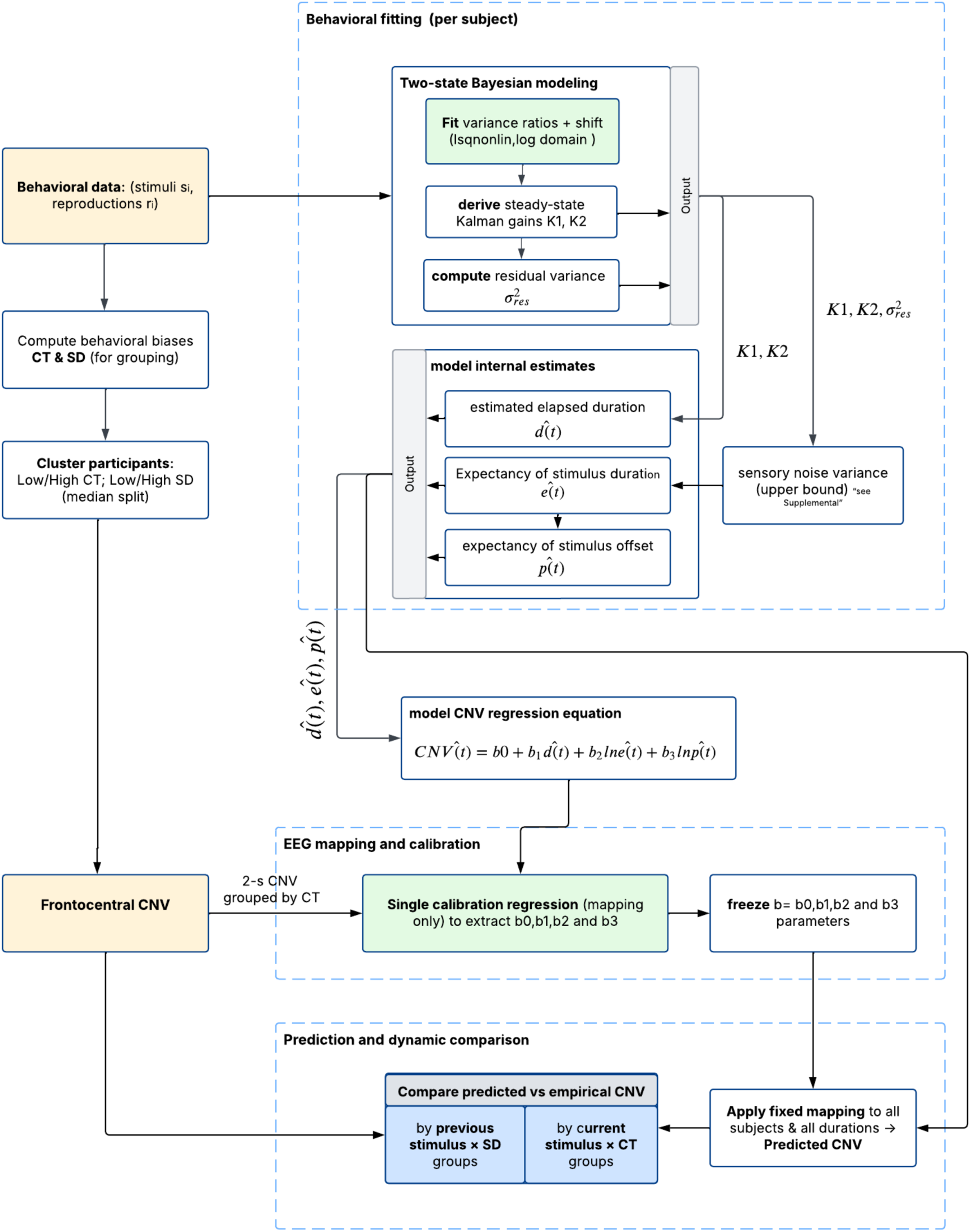
Pipeline linking behavioral two-state Bayesian modeling to CNV predictions. Behavioral data (stimuli *s*_*i*_, reproductions *r*_*i*_) are first used to compute individual perceptual biases central tendency (CT) and sequential dependence (SD). Participants are then median-split into Low/High CT and Low/High SD groups. Next, per subject, behavioral data are fit with a two-state Bayesian model: we fit two free parameters of variance ratios and shift (lsqnonlin) and then derive steady-state gains K1 and K2; the residual variance 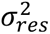 is computed. From 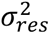 together with K1 and K2, an upper bound on sensory-noise variance 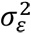 is calculated (see Methods and Supplementary material). Using the fitted model, we compute per-trial three model internal time courses during the production phase: estimated elapsed duration *d̂*(*t*), expectancy density *ê*(*t*) (prior over duration), and its integral, the offset probability *p̂*(*t*). A single calibration maps these variables to the frontocentral CNV (FC1, Cz, FC2) using only 2-s trials grouped by CT in the 0.4–2.1 s window. The predicted 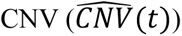 is formed as linear summation of these three components: 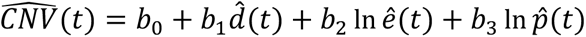. The coefficients *b*_0_ − *b*_3_ are then frozen and applied to all subjects and durations to obtain predicted CNV time courses, which are then compared with empirical CNV by previous-stimulus × SD and current-stimulus × CT groupings (no refitting). Color coding: Yellow = inputs from experimental data; Green = fitting/mapping steps; Blue = final stage/output.

Accordingly, 48 volunteers (22 female, average age 29.5 ± 3.84 years) participated in the experiment. All participants had normal or corrected-to-normal vision. They were naive to the purpose of the experiment and were monetarily compensated for their participation (10 Euro/h). The experiment was approved by the ethics committee of the Brandenburg University of Technology Cottbus-Senftenberg. Informed written consent was obtained prior to the experiment.

### Time interval reproduction task

Participants performed a time interval reproduction task, in which they were asked to perceive and then reproduce presented time intervals. The intervals were indicated by an audio-visual stimulus consisting of a visual signal (white plus sign on a black screen) accompanied by a continuous auditory beep (441 Hz). Only three different durations (1, 1.5 and 2 s) were tested. The stimulus (production phase) was followed by a short break (random duration between 1.0 and 1.125 s). The reproduction phase started with the presentation of a white star sign in the middle of the screen and the auditory beep, and subjects were asked to press the Space key as soon as the stimulus duration was the same as in the production phase. The keypress terminated the audio-visual signal (Fig. 1A).

The experiment was conducted in three blocks, each consisting of 20 randomly ordered trials for each stimulus duration, resulting in a total of 60 trials per block. Throughout the entire experiment, each participant underwent 180 trials across all blocks. Following each block, participants took a break as needed before proceeding to the next block.

Before starting the main experiment, participants did a practice run with 30 trials, after each practice trial but not during the main experiment with EEG, coarse visual feedback with five levels (Glasauer & Shi 2021) was provided which indicated whether the reproduced time interval was within ±5% of the stimulus duration (green middle circle), between 5% and 30% (an orange circle left or right to the middle) (inner orange circle positioned to the left for shorter intervals, and to the right for longer intervals), or larger than ±30% (outer red circles to the far left for shorter intervals and far right for longer intervals). Stimuli were generated and displayed using Matlab (The Mathworks) and Psychtoolbox 3 (Brainard, 1997; Kleiner et al., 2007).

### EEG recording and preprocessing

EEG signals were recorded using a 32-channel LiveAmp amplifier (Brain Products GmbH, Germany) equipped with actiCap wet electrodes. Data were sampled at 500 Hz, and the electrode montage followed an extended version of the international 10 – 20 system, with FCz and Fpz serving as reference and ground electrodes, respectively. Throughout the entire experiment, electrode impedances were maintained below 10 kΩ. An online bandpass filter with cutoff frequencies of 0.01 Hz and 70 Hz was applied, and the data were re-referenced to a common average reference (CAR), calculated from the potentials of all 32 electrodes. Independent Component Analysis (ICA) was performed using EEGLAB software to remove artifacts such as eye blinks, horizontal and vertical eye movements, and muscle activity (Delorme and Makeig, 2004). Prior to ICA, a 0.1 Hz high-pass, zero-phase FIR filter was applied in EEGLAB to reduce slow drifts, improve stationarity, and yield more reliable ICA decomposition while preserving slow CNV dynamics (hence the conservative 0.1 Hz cutoff). ICA decomposition was visually inspected, and artifact-related components were identified based on standard criteria including scalp topography, spectral properties, and component time-courses. Components primarily representing artifacts were manually marked and removed from the data. After ICA artifact removal and further preprocessing, three participants exhibited substantial residual noise or remaining artifact contamination (e.g., movement artifacts, extremely noisy signals), making their EEG data unsuitable for reliable ERP analysis. These participants were therefore excluded from all EEG analyses. Subsequently, further preprocessing, including trial extraction and offline filtering, was conducted using MATLAB’s built-in functions and the FieldTrip toolbox (Oostenveld et al., 2011).

### Behavioral data analysis

Prior to analyzing the reproduced intervals as the behavioral data, we excluded outliers on the basis of the log-transformed reproduction intervals: reproduction intervals that deviated from the mean by more than three times the standard deviation of reproduced intervals were excluded for each subject and stimulus duration separately. This led to exclusion of 0.83% of all trials.

Repeated Measures ANOVA was employed to examine within-subject factors, including the effects of current stimulus duration and previous stimulus duration on reproduction intervals. Interaction effects between sequential dependencies and stimulus durations were also investigated. When necessary, appropriate corrections for sphericity violations were applied. Post hoc comparisons were conducted to investigate interactions between previous stimuli and sequential dependencies, with p-values adjusted for multiple comparisons using Bonferroni’s method. ANOVA analysis was done using JASP (version 0.18; JASP Team 2024) statistical computer software for analysis.

Sequential dependence and central tendency were computed for each participant. We computed central tendency by estimating the slope *m* of the least-squares line fit of current reproduction interval *r*_*i*_ at trial *i* over current stimulus interval *s*_*i*_. The slope (*m*) of this regression reflects how accurately participants reproduced durations, a slope of 1 indicates perfect accuracy, whereas slopes less than 1 indicate regression toward the mean duration (central tendency). Thus, central tendency was quantified as *CT* = 1 − *m*, where higher values indicate stronger central tendency. For the sequential dependence (SD), we calculated the linear regression of current reproduction error *e*_*i*_ = *r*_*i*_ − *s*_*i*_ over previous stimulus duration *s*_*i*−1_ and considered the slope of this regression as sequential dependency. We then used the median values of CT and SD as criterion to partition subjects into two groups with high and low sequential dependence (High SD and Low SD groups) and high and low central tendency (High CT and Low CT groups).

### CNV response analysis

As we were interested in the slow fluctuation of EEG data, particularly CNV responses, a low-pass filter with a cutoff frequency of 7 Hz was applied to the EEG. Subsequently, CNV responses were extracted separately for the production. For the analysis of early CNV responses, all trials were extracted from 0.2 seconds before stimulus onset to 2.5 seconds after. For production-phase ERPs, baseline correction was applied using a −0.2 to 0 s window relative to stimulus onset. For post-production offset ERPs, a symmetrical 100 ms (−50 to 50 ms) window centered on the stimulus offset was used. These intervals were used to normalize each epoch individually. To investigate sensory responses attributed to stimulus offset, a bandpass filter with cutoff frequencies of 1 and 20 Hz was employed to reduce contributions from slow drifts (e.g., CNV), which predominantly occupy lower frequencies below 1 Hz (Kononowicz & van Rijn, 2014). For CNV analysis, we focused on electrode clusters over the frontocentral region (FC1, Cz, FC2) and the parietal region (right parietal: P4, P8; left parietal: P3, P7), given their established roles in temporal processing and attentional mechanisms. The frontocentral region was chosen based on its well-documented involvement for anticipatory processes, temporal expectations, and integration of contextual information during interval timing tasks (Kononowicz & van Rijn, 2014; Ng et al., 2011; Wiener & Kanai, 2016). The right parietal region was explicitly included because of its consistent involvement in subjective duration perception, sequential dependence, and sensory history integration, as repeatedly shown by EEG and fMRI studies (Bueti & Walsh, 2009; Merchant et al., 2013; Hayashi & Ivry, 2020; Akrami et al., 2018). The corresponding left parietal region was included as control. By focusing on these two regions, we aim to capture both the anticipatory mechanisms in the frontocentral area and the higher-level processes such as the integration of sensory history and perceptual biases, represented in the right parietal cortex.

### Bayesian model

We used the Bayesian model published in Glasauer & Shi (2022) to fit behavioral results and to predict possible representations of time perception during the production phase (Fig. 1 for examples). The model estimates the perceived value *z* from the measured value *x* of a stimulus *s*, based on an assumption of how the stimuli are created in the world. This assumption, called the generative model (see Ma et al., 2023), is depicted graphically in Fig. 1B. The generative model assumes that the stimulus *s*_*i*_ presented at trial *i* comes from a normal distribution (*m*_*i*_, *σ*_*s*_) with mean *m*_*i*_ and variance 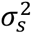. The mean *m*_*i*_ of this stimulus distribution on trial *i* is the the mean *m*_*i*−1_ in the previous trial plus some random change *δ* which is normally distributed with mean 0 as *δ* = (0, *σ*_ẟ_). The evolution of the mean of the stimulus distribution thus corresponds to a time-discrete Wiener process or random walk. In addition, the measured duration *x*_*i*_ is equal to the stimulus *s*_*i*_ corrupted by Gaussian noise so that *x*_*i*_ = *s*_*i*_ + *ε* with *ε* = *N*(0, *σ*_*ε*_), where 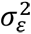 is the variance of sensory noise. The generative model (Fig. 1B) can thus be written as:

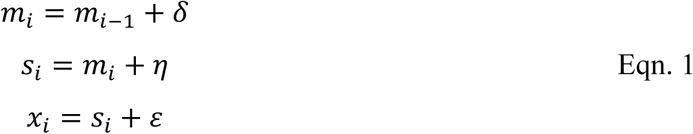

with *η* being normally distributed with *N*(0, *σ*_*s*_). If *δ* = 0 or *σ*_ẟ_ = 0 the model reduces to the boundary case of a static stimulus distribution with fixed mean. If *η* = 0 or *σ*_*s*_ = 0 the model reduces to an iterative model that results in a time-discrete random walk of the stimulus (Glasauer & Shi 2022).

Thus, the generative model covers a continuous range of assumptions about stimulus generation from complete randomness to strong trial-to-trial dependence. Accordingly, the corresponding estimation model can account for different combinations of central tendency and sequential dependence, as shown in Fig. 1.

The estimation model can be expressed as Kalman filter with two free parameters, the ratio between the noise variance 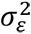 and the variances 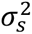 or 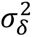. The steady state version of the estimation model is

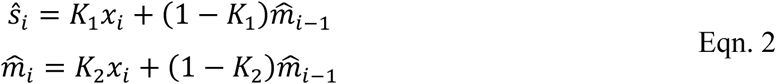

With *ŝ*_*i*_ being the estimate of stimulus duration *s*_*i*_, 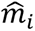 being the estimated mean of the stimulus distribution at trial *i* and *K*_1_ and *K*_2_ the two steady-state Kalman gains. The Kalman gains act as weights and are bounded between 0 and 1, with the additional constraint *K*_1_ ≥ *K*_2_. To account for the scalar property in time perception, the stimuli are log-transformed before applying the estimation model, and the resulting values are back-transformed to yield perceived duration (e.g., Petzschner & Glasauer 2011; Roach et al., 2017). The two Kalman gains are also related to central tendency and sequential dependence (see also Shi et al., 2025): using the equations above without log-transform, it can be shown that for model simulations central tendency is equivalent to 1 − *K*_1_, while sequential dependence is *K*_2_ · (1 − *K*_1_). An additional shift parameter accounts for global under- or overestimation (see Petzschner & Glasauer, 2011; Glasauer & Shi, 2022). The complete Kalman filter model is fit to the raw behavioral data of each participant using Matlab’s non-linear fitting procedure `lsqnonlin` to derive the individual model parameters. Using the individually fitted model, the model’s instantaneous running estimate of elapsed duration during the production phase (Fig. 1C3–E3) and the expectancy of stimulus offset (Fig. 1C4 – E4, see also below) was calculated for each subject and trial.

In addition to the estimated stimulus duration *z*_*i*_, the Kalman filter also estimates the variance of the prior distribution *relative* to the measurement noise distribution. Thus, to determine the instantaneous expectancy of stimulus offset (Fig 1C4 – E4) expressed as probability density of the prior over time, the variance of the measurement noise 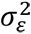 must be estimated. The variance of the residuals 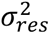, i.e., of the difference between model fit and behavioral duration, cannot be used directly, because the effect of measurement noise on the responses is minimized by the estimation process. However, assuming that the residual variance 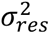 is caused only by optimized effect of measurement noise (and not by additional execution noise during reproduction), an upper bound for the sensory noise variance 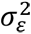 can be determined (see Supplemental Material). The sensory noise variance estimated in this way depends not only on the residual variance but also on the steady-state Kalman gain and differs for low and high sequential dependence: for the static boundary case *K*_2_ = 0 (sequential dependence zero) the sensory variance is 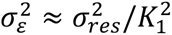, for the random walk boundary case *K*_2_ = *K*_1_ (sequential dependence maximal) it is 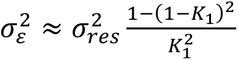. The estimate of the noise variance is determined for each individual and used for prediction of the prior distribution and the instantaneous expectancy (see Fig 1C4 – E4). For example, in Fig 1E1 – E4 estimated elapsed time is close to physically elapsed time, i.e., almost no central tendency, not just for behavioral responses (Fig 1E1), but also for estimated time during production (Fig 1E3). Accordingly, the expectancy for stimulus duration is largest approximately for the mean stimulus duration of 1.5 s (Fig 1E4). In addition to the expectancy expressed as prior probability density (Fig. 1C4 – E4), the integral of the prior density, the cumulative prior probability distribution, can serve as expectation for stimulus offset, because it represents the probability that the stimulus ends right now. This probability increases monotonically with time from zero to one (not shown) and is also used for modeling the CNV.

### Modeling the CNV

To simulate the time course of the CNV, we used three model variables (or components) determined over time: the estimated elapsed duration since stimulus onset *d̂*(*t*) (Fig. 1C3 – E3), the instantaneous expectancy of stimulus duration *ê*(*t*) expressed as prior probability density (Fig. 1C4-E4), and its integral, the probability for stimulus offset *p̂*(*t*). The estimated elapsed duration *d̂*(*t*) for trial *i* was computed using Eqn. 2 using the respective Kalman gains *K*_1,*i*_ and *K*_2,*i*_ for that trial and participant and elapsed time *x*(*t*) = *t* as input. Expectancy of stimulus duration *ê*(*t*) was computed using the relative prior variance of *d̂*(*t*) resulting from the Kalman filter and the estimated measurement variance (see above, and Supplemental Material), and *p̂*(*t*) was calculated as the integral of *ê*(*t*). Note that all three variable time courses were calculated for each single stimulus and for each participant, so that for each measured CNV there were also three predicted time courses of the model variables.

We then assumed that the average CNV might represent a linear combination of the three variables (averaged over stimuli and participants) with the probabilities represented as log probability, as suggested previously for representing probabilities in the brain (Haefner et al., 2024):

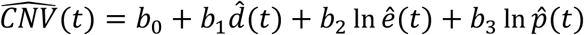

with *b*_0_ being an offset term and *b*_1,2,3_ three scaling factors. This model estimate 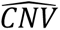 was fit to the average CNV, grouped by central tendency, of the longest stimulus (2 s) in the time window 0.4 – 2.1 s by linear regression. The resulting offset and three scaling factors were then applied for all further CNV simulations. Fig. 2 summarizes the modeling pipeline from behavioral fitting of the two-state model to CNV predictions and comparisons.

## Code Accessibility

The MATLAB code and behavioral data used for Bayesian modeling in this study are freely available online at: https://gin.g-node.org/sglasauer/CNV_prediction. EEGLAB (Delorme & Makeig, 2004) and the FieldTrip toolbox (Oostenveld et al., 2011) were used for preprocessing and data analysis, and the PsychToolbox (Brainard, 1997) was used for the stimulus presentation paradigm.

## Results

### Behavioral Results

Fig. 3A shows the mean reproduction errors of all subjects plotted over stimulus duration. The reproduced durations exhibit considerable central tendency, that is, high durations (2s) tend to be underestimated, while lower durations (1s) are overestimated. In Fig. 3B and 3C the reproduction errors are plotted separately for each previous stimulus duration and the two groups of subjects separated by sequential dependence. Fig. 3B shows the errors for the group of subjects with low sequential dependence (low SD), Fig. 3C for those with high sequential dependence (high SD). Repeated measures ANOVA on reproduction errors with between-subject factor *SD group* (high and low SD) and within-subject factors *current stimulus* and *previous stimulus* (3 levels each: 1 s, 1.5 s, and 2 s) shows a significant main effect for the factor *current stimulus* (F(2,92) = 362.137, p < 0.001, η^2^ = 0.491), but no interaction between *current stimulus* and *SD group* (F(2,92) = 2.32, p = 0.1, η^2^ = 0.003). Thus, the effect of the current stimulus on reproduced intervals did not significantly differ between subjects with high and low sequential dependencies. The factor *previous stimulus* showed a significant main effect (F(2,92) = 43.65, p < 0.001, η^2^ = 0.014). As expected from the between-subjects grouping criterion, and thus not being regarded as result, the interaction between the factor *previous stimulus* and *SD group* was significant (F(2,92) = 24.706, p <0.001, η^2^ = 0.008). Fig. 3D shows central tendency versus sequential dependence for both subject groups. As expected from the missing interaction between *current stimulus* and *SD group*, we did not find significant differences between the central tendencies of the two groups (*t*(46) = 1.75, *p* = 0.086).

**Figure 3.**
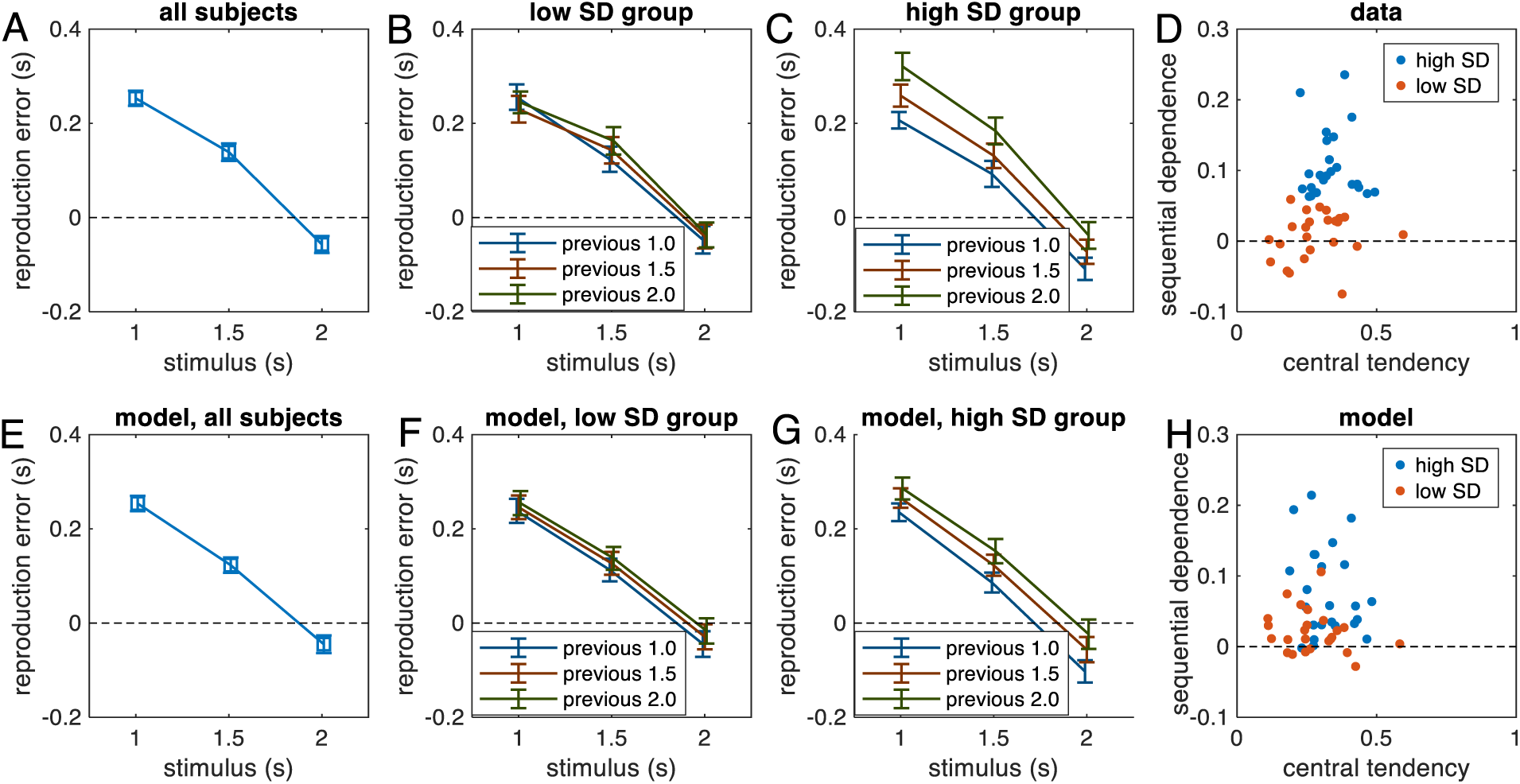
Results of behavioral data analysis and model simulation. (error bars indicate standard error of the mean): **A.** Mean reproduction error for all subjects (n = 48) plotted over stimulus interval. The dependence of reproduction error on stimulus duration shows a strong overall central tendency. **B.** Mean reproduction error plotted over current stimulus duration separately for each previous stimulus duration for subjects with low sequential dependency (n = 23). **C.** As in B, but for subjects with high sequential dependency (n = 22). **D.** Sequential dependence (SD) plotted over central tendency for subjects with high (blue dots) and low (red dots) sequential dependence. **E – H.** As in A – D, but model simulations, the grouping in F – H is the same as in B – D. For simulation, the model was fitted to the raw responses of each participant separately and the model responses were then treated the same way as the participants’ responses. **H.** The simulated sequential dependence of the model simulations for the two groups shows more overlap than the data in D.

### Modeling the behavioral responses

We applied the two-state model (Glasauer & Shi 2022), which was developed to explain behavioral responses in magnitude reproduction experiments within a Bayesian framework, to the behavioral data. The model (see Methods, 3 free parameters: assumed relative variance of the stimulus distribution, assumed relative variance of random changes of the mean of the stimulus distribution, and an overall bias) was fit to each participant separately and the model responses were then treated and analyzed the same way as the behavioral responses. Fig. 3E – H depicts the average model responses. It should be noted that the good fit of the model to the data is not a trivial result, since in the present experiment only three different stimuli were tested, while the model was developed for stimuli from a quasi-continuous range. To determine the goodness of fit of the model and whether all parameters are required, we also fitted the model without one of the variance parameters, so that the two comparison models (see also Glasauer & Shi 2022), each having just two parameters, corresponded to a static Bayesian model (with fixed prior distribution) and a simple iterative model (in which the percept of the current trial is the prior for the next trial). The steady-state version of the latter corresponds the internal reference model (Dyjas et al. 2012; Bausenhart et al. 2014). The AIC values showed a significant effect of “*model*” (F(2,92) = 83.767; p < 0.001, η^2^= 0.010) with best values for the two-state model (average for each model: two state −544, static −543, iterative −522) and a significant interaction between “*model*” and “*SD group*” (F(2,92) = 10.75; p < 0.001, η^2^ = 0.001), but no main effect of *SD group*. The *model* x *SD group* interaction is expected, because for the low SD group, the static model can be sufficient, while for the high SD group the iterative model could suffice. However, taken together, our analysis confirms that the two-state model on average provided a better fit than both simpler models.

We also evaluated the fitted model parameters with relation to the subject groups. From the relation between Kalman gains and biases (see Methods), it is expected that *K*_1_ is related to central tendency, and *K*_2_ to sequential dependence. Indeed, *K*_2_(0.121 ± 0.144, mean ± SD) showed a significant difference between subjects with *low SD* and *high SD* (t(43) = −2.10, p = 0.042), but not between subjects with *low* and *high* central tendency (t(43) = 0.18, p = 0.86). Conversely, *K*_1_ (0.649 ± 0.095) was not significantly different for the *SD groups* (t(43) = 1.28, p = 0.21), but highly significantly different for subjects with *low* and *high* central tendency (t(43) = 7.34, p < 0.0001).

From the Kalman gain parameters *K*_1_ and *K*_2_ of the individual models and the individual residual variance values we also determined the variance of measurement noise for each participant (see Methods), which is used for predicting representations of model variables during production for comparison with the CNV (see below). Expressed as standard deviation of the measurement noise distribution the average over all participants was 0.35 ± 0.12 s (mean ± SD). There was no difference between subjects with low SD and high SD (t(43) = −0.55, p = 0.58). However, when grouped according to central tendency, there was a highly significant difference with higher sensory noise for higher central tendency (t(43) = −2.8, p = 0.0076). This result is expected, first because low measurement noise should cause higher reliance on stimulus magnitude, which in turn causes low central tendency, and second because the variance is estimated using the Kalman gain *K*_1_, which is approximately inversely proportional to central tendency (see Methods).

### Neural correlates of sequential dependence and central tendency

We first evaluated averaged CNV responses in the frontocentral region (FC1, Cz, FC2) measured during the early production phase (time window 0.8 – 1.2 s). A repeated measures ANOVA with main factor of *current stimulus* was conducted for all subjects to evaluate the CNV amplitude across different stimulus durations (1s, 1.5s and 2s). As expected from the randomized stimulus presentation, there were no significant differences between early CNVs depending on *current stimulus* duration (F(2,88) = 0.752, p = 0.474, η^2^ = 0.017). It is expected, because in the early production phase participants cannot know how long the current stimulus will last.

We then evaluated whether *previous stimuli* influence neural responses during the early production phase in the frontocentral region. A repeated measures ANOVA (n=45, see Fig. 4A1 and 4C1) confirmed a significant dependence of CNV amplitude (time window 0.8 – 1.2 s) on *previous stimulus* (F(2,88) = 4.681, p = 0.012, η^2^ = 0.096) confirming a recent study (Damsma et al. 2021).

**Figure 4:**
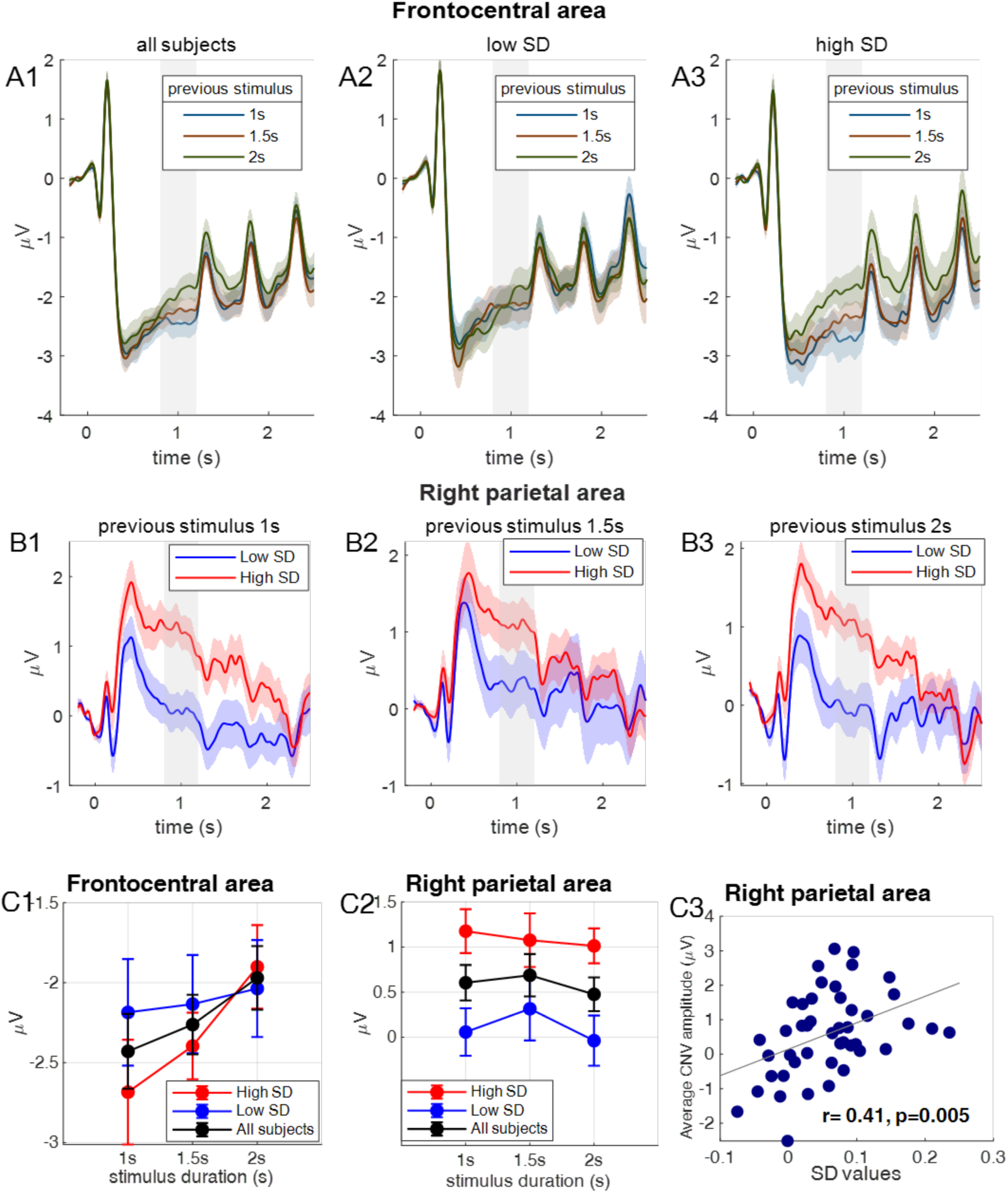
Effect of previous stimulus on CNV amplitude in the frontocentral and right parietal lobe: **A1.** Grand average production phase response in frontocentral area for trials where the previous stimulus duration is 1 s, 1.5 s or 2 s for all subjects (n=45). **A2.** Same for subjects with *low* sequential dependence (n = 23) and **A3.** for subjects with *high* sequential dependence (n = 22). **B1.** Grand average production phase response in the right parietal area for stimulus duration 1 s, **B2**. 1.5 s, and **B3**. 2 s. The gray bar denotes the time window for CNV amplitude quantification**. C1.** Frontocentral CNV amplitude plotted over previous stimulus duration, blue: group with low sequential dependence (*low SD*), red: group with high sequential dependence (*high SD*), black: all subjects. **C2.** right parietal CNV plotted over previous stimulus duration. Colors as in C1. Error bars denote standard error of the mean. **C3.** Correlation between individual sequential dependence values and average right parietal CNV amplitude (0.8–1.2 s). Each dot represents one subject (n = 45); a significant positive correlation was observed (r = 0.41, p = 0.0047).

The difference in individual sequential dependence observed in the *high SD* group vs. *low SD* group of our subjects further suggests that neural responses during the production phase might be influenced differentially by previous stimulus duration. We therefore conducted a repeated measures ANOVA with within-subject factor *previous stimulus* (1s, 1.5s, 2s) and between-subject factor SD *group* (*high SD* vs. *low SD*). As expected, a significant effect of the factor *previous stimulus* was found (F(2,86) = 4.96, p = 0.009, η^2^= 0.019). However, neither the factor *group* (F(1,43) = 0.303, p = 0.585, η^2^ = 0.006) nor the interaction *group* x *stimulus* (F(2,86) = 1.154, p = 0.107, η^2^ = 0.009) became significant. The average CNV responses for both groups are shown in Fig. 4A2 and 4A3. Thus, even though the influence of previous stimuli on behavior was significantly different between groups, their influence on frontocentral CNV responses in the time window 0.8 – 1.2 s was not. We furthermore observed pronounced differences between neural activity patterns in the right parietal lobe electrode cluster (P4 and P8) for individuals with high compared to low sequential dependence (Fig 4B1 – B3, 4C2). A repeated measures ANOVA was conducted to investigate the effects of *previous stimulus* duration and *SD group* on CNV amplitude in the time window 0.8 –1.2s. This time window was selected to determine the CNV amplitude independently of stimulus duration, i.e., remaining below the earliest reaction to stimulus offset, but starting late to capture differences between groups. A significant main effect of *group* was found (F(1,43) = 8.32, p = 0.006, η^2^ = 0.126). As shown in Fig. 4C2, individuals with *high* serial dependence exhibited enhanced neural responses in the right parietal lobe compared to those with *low* sequential dependence. The results showed no significant main effect of *previous stimulus* duration on the neural response (F(2,86) = 0.468, p = 0.514, η^2^ = 0.004), and no interaction between *previous stimulus* duration and subject *group* (F(2,68) = 0.536, p = 0.628, η^2^ = 0.003). No significant effects were found for left parietal electrode clusters (see Supplemental Material Fig. S1).

We then evaluated whether the found *SD group* difference would also apply to individual SD values. To do so, we correlated individual right-parietal CNV amplitude and SD values. The correlation became significant (n = 45, r = 0.414, p = 0.0047), suggesting that right-parietal CNV values may represent a neural correlate of individual sequential dependence (Fig. 4C3).

### CNV amplitude in individuals with different central tendencies

We investigated whether frontocentral CNV amplitude differs between subjects with different levels of central tendency. Figure 5 shows the average ERP during the production phase for all subjects (Fig. 5A) and separately for those with low (Fig. 5B) and high central tendency (Fig. 5C). A repeated measures ANOVA was conducted to examine the effects of *current stimulus* duration and central tendency *group* on CNV amplitude, calculated as the mean value within the 0.8–1.2s time window. A significant main effect of *group* was found, with subjects exhibiting lower central tendency values showing higher CNV amplitudes (F(1,43) = 5.085, p = 0.029, η² = 0.083; see Fig. 5D). However, as expected, there was no significant effect for the factor *current stimulus* (F(2,86) = 0.73, p = 0.48, η² = 0.004), nor was there a significant interaction (F(2,86) = 0.22, p = 0.818, η² = 0.001).

**Figure 5:**
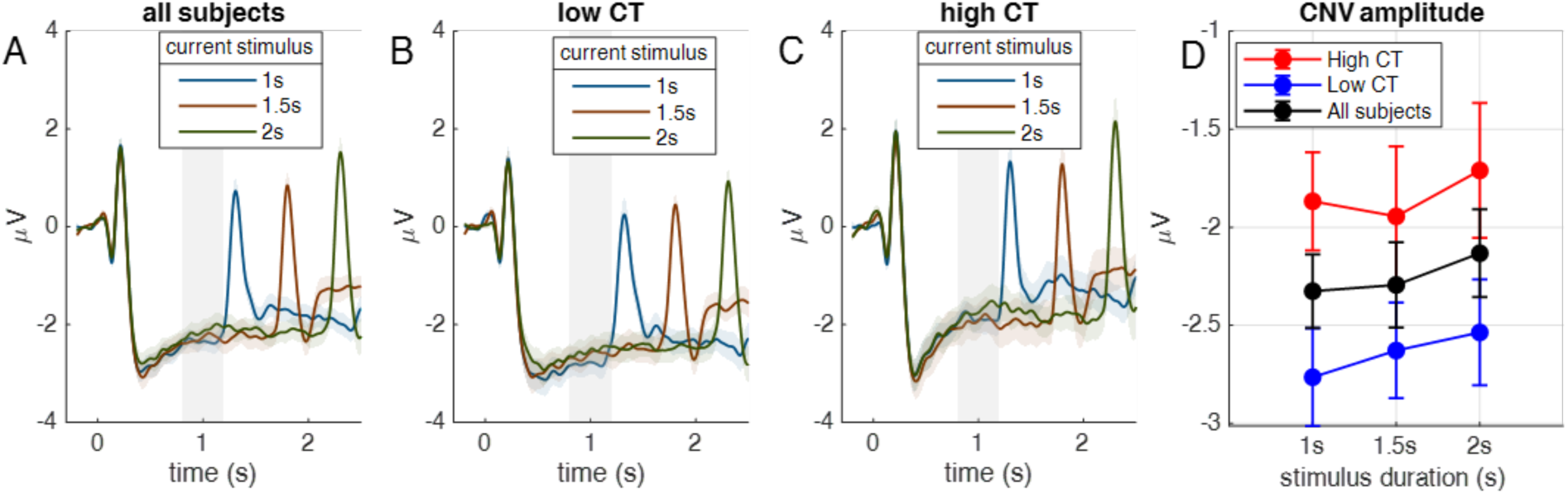
Effect of the current stimulus on CNV amplitude in the frontocentral lobe, with subjects grouped by central tendency: Grand average response for trials where the current stimulus durations are 1s, 1.5s, and 2s for **A**. all subjects **B.** Subjects with low central tendency and **C.** Subjects with high central tendencies. Gray bars show time window for CNV amplitude quantification (0.8 – 1.2s) **D.** CNV amplitude plotted over current stimulus duration. Blue: group with low central tendency (*low* CT, n = 23), red: group with high central tendency (*high* CT, n = 22, black: all subjects (n = 45). Error bars denote standard error of the mean.

Again, we evaluated whether this group difference would also apply to individual CT values. However, the correlation between individual frontocentral CNV and CT values did not reach significance (n = 45, r = 0.18, p = 0.21).

### Linking Bayesian estimation of elapsed time to neural activity during stimulus production

We next asked whether the predictions of our Bayesian model for estimating elapsed duration during the production phase shown in Fig. 1 could be matched to the frontocentral CNV. To do so, we used the fitted individual model parameters (see above) and simulated the individual models for all participants and all trials to compute a running estimate of elapsed time for each stimulus during the production phase, so that for every single CNV there was one corresponding simulation (see also Methods). We assumed that the CNV might be a linear combination of three components of the model: 1) estimated elapsed time since stimulus onset (Fig. 1C3 – E3), 2) estimated expectancy of stimulus offset as log prior probability density (Fig. 1C4 – E4), and 3) the probability for stimulus offset (the log cumulative prior density, not shown, integral of the curves in Fig. 1C4 – E4). We first evaluated whether the average CNV for the longest stimulus duration 2 s (green curve in Fig. 5A) could be approximated by one of the averaged model components. To make the model components comparable to the CNV, we determined a scaling factor for each possible predictor (model component) of the CNV and an offset by fitting the grand average model prediction to the grand average CNV of stimulus duration 2.0 (time window 0.4 – 2.1 s) by linear regression. Fig. 6 shows the result of fitting single model components to the CNV. The best result was achieved by prior expectancy with a coefficient of determination R^2^ = 93.9%.

**Figure 6:**
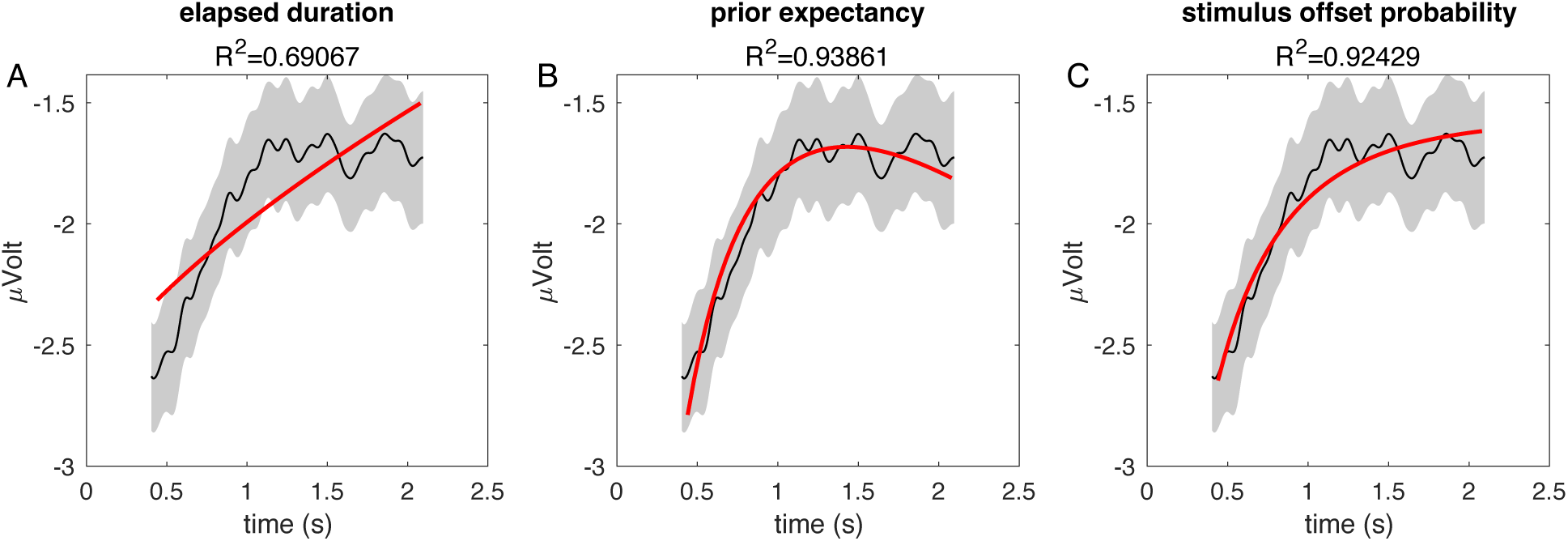
Frontocentral CNVs for 2 s production phase (black, same data as in Fig. 5A) and simulated model CNV (red) for different single variables. (**A**: estimated elapsed duration, **B**: prior expectancy of duration, **C**: stimulus offset probability; see text and Fig. 1). The CNV is averaged over stimuli and participants (n = 45). Shaded area shows standard error of the mean. The model prediction is also averaged over stimuli and participants (n = 45). The averaged model variable is fitted to the average CNV (time window 0.4s – 2.1s as depicted) by linear regression, the corresponding R^2^ values are shown on top.

While this result was encouraging and worked well also for the grand average CNVs of the other two stimulus durations (not shown), it was not able to explain the differences between CNVs of different CT groups (Fig. 5B vs. 5C). We therefore chose to fit linear combinations of all three model components to the average 2 s CNV of both groups instead to the grand average. The combination of all three predictor components achieved the best AIC (coefficient of determination R^2^ = 60.0%; AIC = 29.6), the second-best AIC (57.1) resulted from including estimation elapsed duration (Fig. 1C3 – E3) and prior expectancy of stimulus duration (Fig. 1C4 – E4). The data and the corresponding fitted model predictions are shown in Fig. 7F.

**Figure 7:**
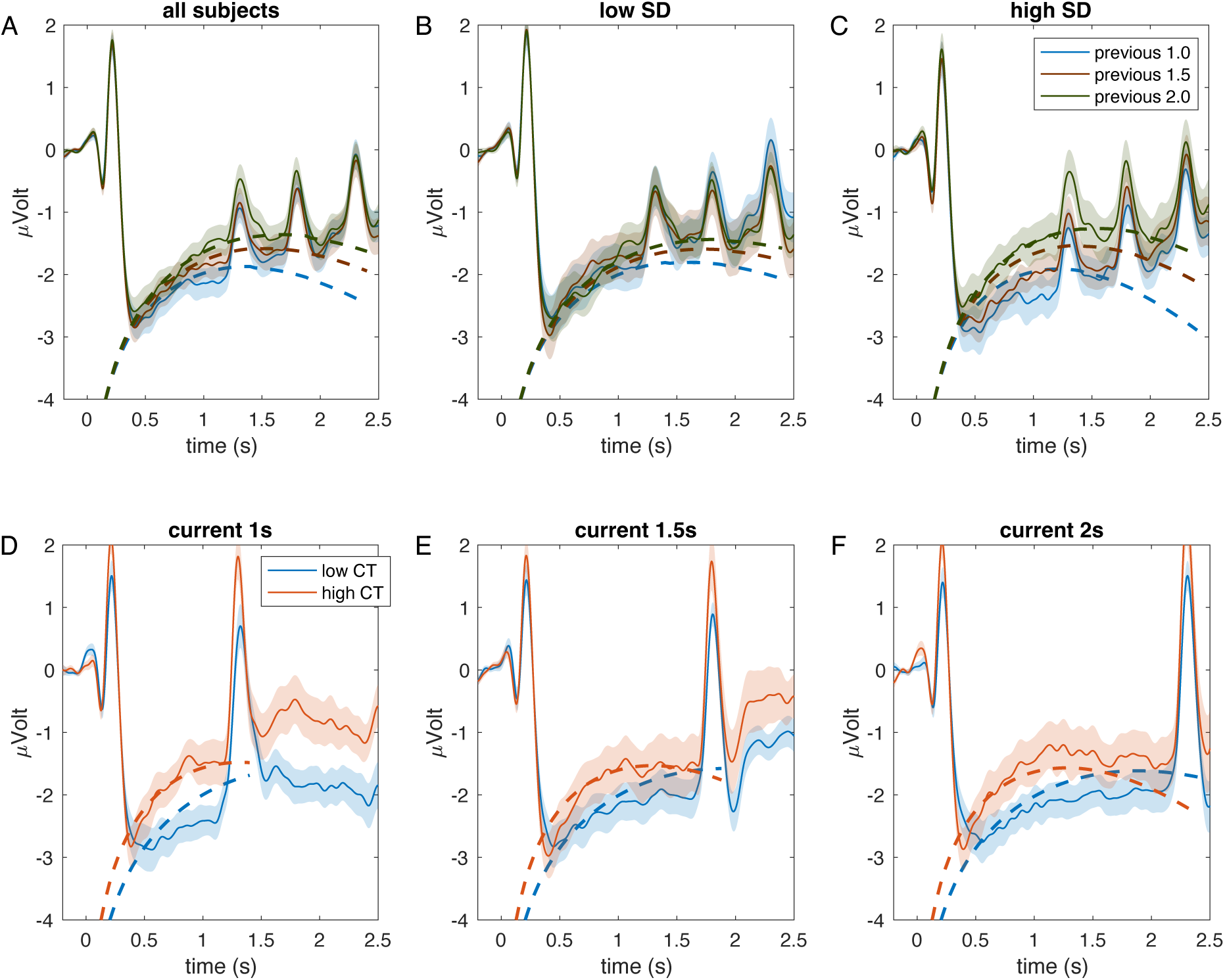
Frontocentral CNVs during production (solid) and predicted model CNV (dashed) for grouping by previous stimulus duration and sequential dependence (A – C, same data as in Fig. 4 A1-A3, colors denote previous stimulus) and grouping by current stimulus duration and central tendency (**D – F,** same data as in Fig. 5 B-C). **A**: all subjects (n = 45), **B,C**: groups with low (n = 23) and high SD (n = 22), **D – F**: blue: groups with low CT (n = 23), red: group with high CT (n = 22) for all stimulus durations. All model simulations (dashed) are, as the CNV data, averages of individual simulations of each trial and each subject. Shaded areas denote standard error of the mean. Note that only for 7F the model predictions derived from the behavioral data have been fitted to the CNV by linear regression (see text, R^2^ = 0.60). Model simulations in 7A – E are predictions using the same regression parameters as in 7F.

The obtained three scaling factors and one offset for the best linear combination were then used to scale and shift all other model simulations in which we evaluated whether the predicted CNVs would show the same differences as the data if grouped according to central tendency or sequential dependence. To quantify whether the model predictions provide a good match to the CNV data, we calculated the correlation coefficients between each model CNV and data CNV shown in Fig. 7B - F.

Fig. 7A – C shows data together with predicted CNVs grouped by previous stimulus duration for all subjects (Fig. 7A) as well as the low SD and high SD groups (Figs. 7B and 7C). In Fig. 7A (data as in Fig. 4A1), the model simulation shows a similar tendency as the data for all subjects together: the CNV is most positive for the longest previous duration (2.0 s, green), and most negative for the smallest previous duration (1.0 s, blue). For the low SD group, differences are smaller (Fig. 7B), while they become more pronounced for the high SD group (Fig. 7C). The correlation coefficients between predicted and experimental CNV grouped by SD and previous stimulus was calculated for the time window 0.4 – 1.1 s and ranged from 0.926 to 0.987 (all p < 1e-7) with the lowest value for high SD and previous stimulus duration 1s (Fig. 7C blue lines) and the best correlation for high SD with 2 s previous duration (Fig. 7C red lines).

It should be emphasized again that the *only* fit of CNV simulation to CNV data was the linear regression performed on the average CNV for 2.0 s of the two CT groups shown in Fig. 7F. The corresponding correlation coefficients in the time window 0.4s - 2.1s for this case were 0.923 for low CT and 0.930 for high CT (p < 1e-17).

In Fig. 7D –F data and model simulations are grouped by current stimulus duration (as in Fig. 6A – C) and by low central tendency (low CT, blue) vs. high central tendency (high CT, red). As for the grouping in Fig. 7A – C, the differences between groups are similar between model simulation and data not only in direction, i.e., more positive CNV for high CT than for low CT, but also in temporal lag with high CT leading the CNV of low CT. Fig. 7F shows the data to which the model components were fitted and the corresponding simulation. For the predictions in Fig. 7D (current duration 1s) the correlation coefficients (window 0.4 – 1.1 s) were 0.873 (low CT) and 0.984 (high CT). The predictions in Fig 7E (current duration 1.5s) correlated (window 0.4 – 1.6s) with 0.923 (low CT) and 0.982 (high CT). For all correlation coefficients for model-data comparison in 7D and 7E, p < 1e-6.

Despite these excellent correlations between data and predictions, it should be noted that there is an abundance of model variants that might provide similar fits but that we did not test, because more experimental conditions would be required to reliably distinguish these model variants. We hope that the match shown here between model and data helps to suggest further possible explanations for the constituents of the CNV.

We then performed the same analysis for the right parietal CNV. While the fit to the average CNV of the 2.0 s stimulus production phase was reasonably successful only when all three components were combined (coefficient of determination R^2^ = 87.5%), the results shown in Fig. 4B1 – B3 could not be replicated qualitatively by the model simulations. We suppose that the group differences exhibited by the parietal CNV are not due to variables of the internal estimation process but might rather be related to the parameters of this process, in this case being related to the extent of how much the previous trial is used as prior information for the current stimulus duration. In modeling terms, this would be the variance 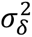 that describes how much the stimulus distribution is changing from trial to trial, or the gain *K*_2_, which is closely related.

### Postproduction Offset ERP: Perceptual Biases and Neural Correlates

Fig. 8A1 depicts the grand average response of all subjects resulting from the offset of stimuli at the end of the production phase in the frontocentral region. In this phase two main peaks, P2 (time window 0.25–0.32 s) and N2 amplitudes (time window: 0.45–0.52 s), were analysed. A repeated measures ANOVA revealed a significant effect for the main factor *current stimulus* on P2 amplitude among all subjects (F(2,88) = 22.15, p < 0.001, η^2^ = 0.33). Like P2, the ANOVA of N2 amplitude showed a significant effect for main factor of *current stimulus* for all subjects (F(2,88)=9.86, p<0.001, η^2^=0.183). We further evaluated whether P2 and N2 amplitudes differ for the two sequential dependency *groups* (Fig. 8A2 – A3). As expected, P2 amplitude showed a significant main effect of *current stimulus* (F(2,86) = 21.79, p < 0.001, η^2^ = 0.092) but no effect of *group* (F(1,43) = 0.273, p = 0.60, η^2^ = 0.005 see Fig. 8D1) and no interaction between *group* and *current stimulus* (F(2,86) = 0.58, p = 0.56, η^2^ = 0.003). N2 amplitude also depended on *current stimulus* (F(2,86) = 9.703, p <0.001, η^2^ = 0.094), but marginally significantly differed among the *high SD* and *low SD groups* F(1,43) = 3.79, p = 0.05, η^2^ = 0.039, Fig. 8D2) with no interaction between *current stimulus* and *group* F(2,86) = 0.79, p= 0.45, η^2^= 0.008).

**Figure 8.**
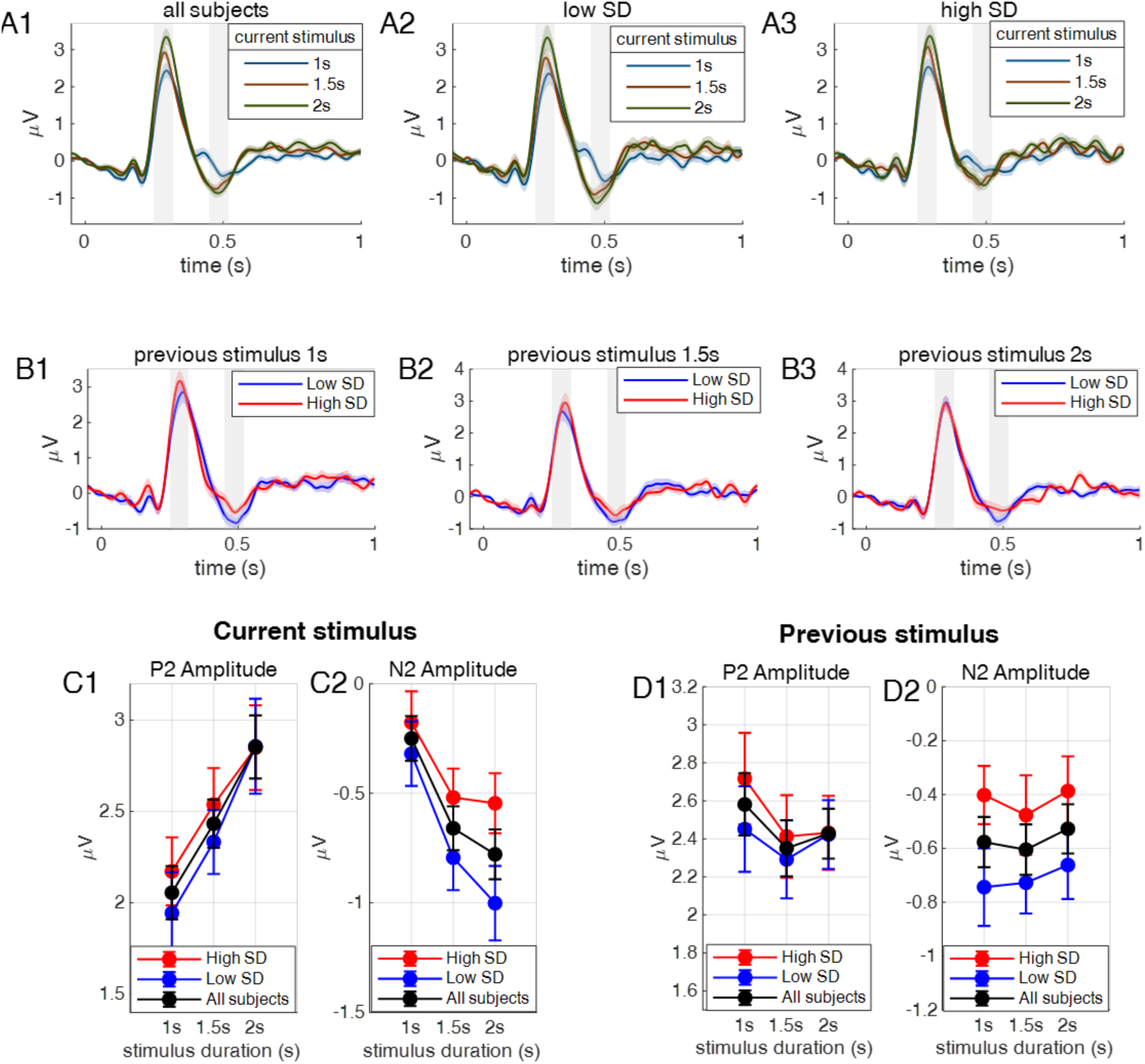
Frontocentral postproduction offset ERP. **A1**: Grand average response of postproduction offset ERP for all subjects (n = 45). **A2**. For low SD group (n = 23). **A3**. For high SD group (n = 22). **B1 – B3**. Postproduction ERP plotted separately for each previous trial stimulus duration (**B1**: 1s, **B2:** 1.5s, **B3**: 2s) for subjects with low SD (blue) and subjects with high SD (red). **C1 – C2.** Postproduction P2 and N2 amplitudes plotted over *previous stimulus* duration for subjects with low SD (blue), high SD (red), and all subjects (black). **D1 – D2.** Postproduction P2 and N2 amplitudes plotted over *current stimulus* duration for subjects with low SD (blue), high SD (red), and all subjects (black). Error bars in C1 – D2 denote standard error of the mean.

We also evaluated whether P2 and N2 amplitudes were affected by the *previous stimulus* duration (Fig. 8B1–B3). A repeated measures ANOVA on P2 amplitude revealed no main effect of *previous stimulus* (F(2,86) = 2.3, p = 0.1, η^2^ = 0.01, see Fig. 8C1), no effect of *group* (F(1,43)=0.23, p=0.62, η^2^=0.004) and no interaction (F(2,86) = 0.68, p = 0.5, η^2^ = 0.003). For N2, the main factor *previous stimulus* was not significant (F(2,86) = 0.389, p = 0.679, η^2^ = 0.003), and N2 amplitude didn’t differed significantly between the *groups* (F(1,43) = 3.669, p = 0.062, η^2^= 0.055, Fig. 8C2) with no interaction between *previous stimulus* and *group* (F (2,86) = 0.141, p = 0.869, η^2^=0.0009).

We then evaluated if the P2 and N2 amplitudes differed between the *groups* of individuals with high and low central tendency (Fig. 9). A repeated measures ANOVA for P2 amplitude revealed a significant difference between central tendency *groups* (F(1,43) = 4.5, p = 0.039, η^2^ = 0.068). As expected from the grouping, the main effect of *current stimulus* duration was also significant (F(2,86) = 21.92, p < 0.001, η^2^ = 0.092) with no interaction (F(2,86) = 0.827, p = 0.481, η^2^ = 0.003). For N2 amplitude, the main effect of *current stimulus* duration was also significant (F(2,86) = 9.8, p < 0.001, η^2^ = 0.093). However, there was no significant difference for the *group* factor (F(1,43) = 7.4×10^−5^, p = 0.993, η^2^ = 8.4×10^−7^) nor for its interaction with *current stimulus* (F(2,86) = 1.35, p = 0.26, η^2^ = 0.013).

**Figure 9.**
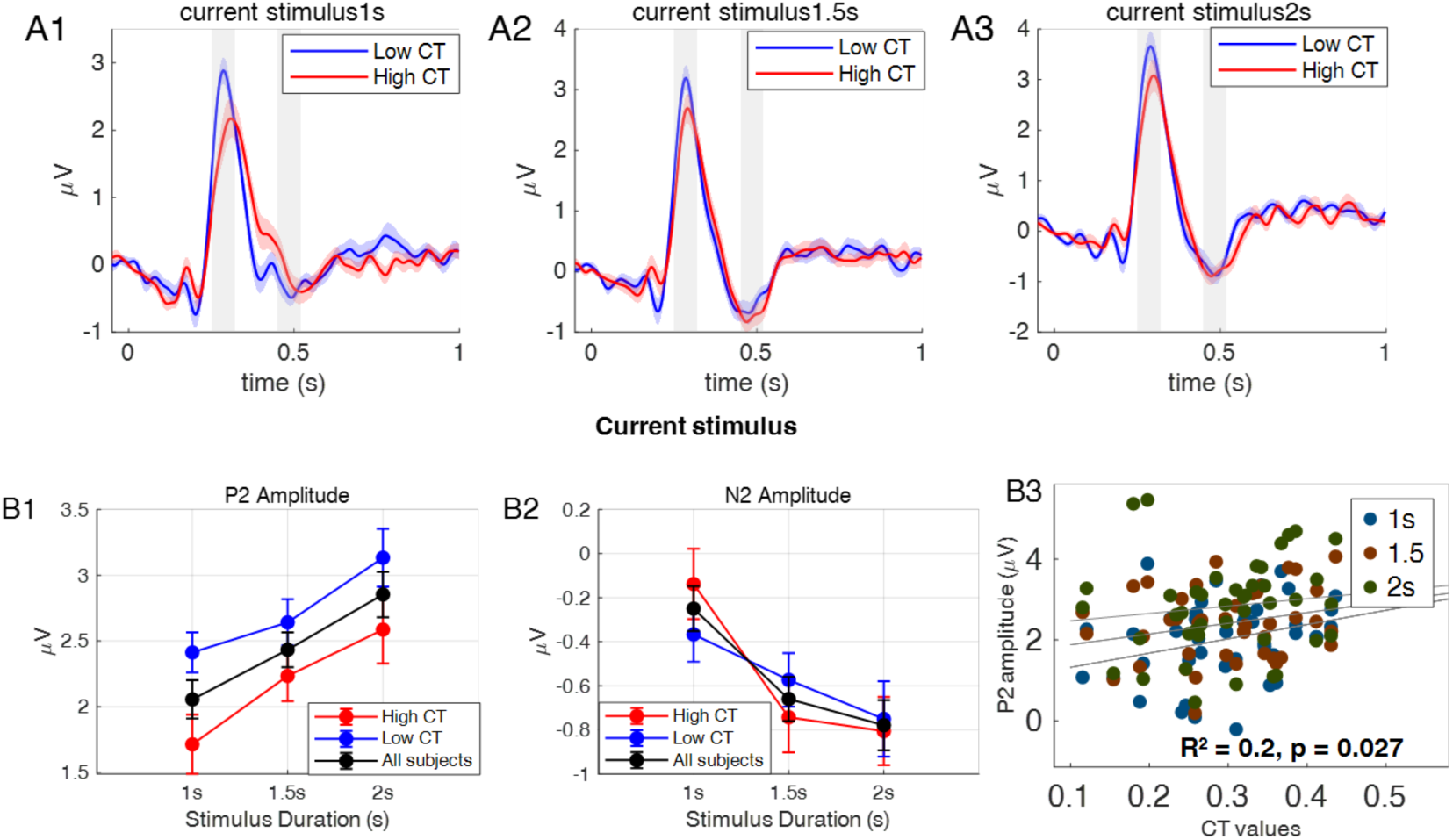
Frontocentral postproduction offset ERP separated for subjects with low (blue) and high (red) CT. **A1**: Grand average response of post-production ERP for 1s. **A2**: 1.5s, **A3**: 2s. Gray bars indicate temporal windows for evaluation of P2 and N2 amplitudes. **B1:** postproduction P2 amplitude depends significantly on current stimulus duration and on subject group (low CT blue, n = 23; high CT red, n = 22; all subjects black, n = 45)**. B2**: postproduction N2 amplitude plotted as in B1. Error bars in B1 and B2 indicate standard error of the mean. **B3**: Scatterplot of individual central tendency values plotted against average P2 amplitudes in the frontocentral region, collapsed across stimulus durations (1 s = blue, 1.5 s = green, 2 s = brown). Each dot represents one participant. Regression analysis indicated a significant association (F(3,41) = 3.42, p = 0.02, R² = 0.20).

Again, we evaluated whether the *CT group* difference for P2 would apply on an individual level. Since P2 also shows a strong dependence on stimulus duration, we chose a regression approach by predicting individual CT values using the average individual P2 amplitudes for all three durations. The regression became significant (F(3,41) = 3.42, p = 0.02, R² = 0.20), which shows that P2 amplitude is related to central tendency bias on an individual level (Fig. 9B3). In contrast to the frontocentral region, postproduction ERPs of the parietal region did not reveal any significant effect regarding sequential and central tendency biases (See Fig. S2 and S3).

## Discussion

Our behavioral results confirm the existence of large individual differences in sequential dependence and central tendency in magnitude reproduction (e.g., Glasauer & Shi 2022). Sequential dependence quantifies to which extent the current reproduction depends on the previous stimulus. Central tendency quantifies overestimation of small stimuli and underestimation of large ones and depends how much the reproduction relies on prior knowledge. Here we show for the first time that these individual differences in perceptual biases are reflected in differences in neural responses measured by EEG, and that both perceptual biases have neural correlates at the individual level. The CNV amplitude in the right parietal area during time perception correlated significantly with the individual behavioral sequential dependence: subjects with high sequential dependence showed a more positive CNV amplitude and vice versa. The individual central tendency bias was significantly related to the postproduction P2 amplitude at stimulus offset with higher P2 peaks for participants with lower central tendency and vice versa.

Furthermore, we confirmed previous results (Damsma et al., 2021) showing that frontocentral CNV amplitude during the production phase is affected by previous stimulus duration (Fig. 4A1 – 4A3), i.e., by the immediate temporal context. Surprisingly, this dependence of CNV on previous stimulus magnitude was not modulated significantly by the behaviorally found sequential dependence (Fig. 4C1). Instead, frontocentral CNV showed a significant relation with central tendency, with more positive CNV for participants with high central tendency and vice versa (cf. Fig. 5).

The right parietal CNV was, in contrast to the frontocentral CNV, not related to previous stimulus duration but showed a clear relationship to sequential dependence (Fig. 4C2). The right inferior parietal cortex has previously been shown to be involved in the encoding of time intervals and the management of attentional readiness and resource allocation (Mohl, 1991; Rao et al., 2001; Bueti and Walsh, 2009; Matthews and Meck, 2016) as well as in subjective duration perception influenced by sensory history (Hayashi & Ivry, 2020). Our results thus suggest that the right parietal CNV amplitude represents, akin to attentional effort, how strongly the percept of the previous stimulus interval affects the estimate of elapsed time in the current interval.

To better understand possible reasons for these dependencies, we used our previously published Bayesian model of time perception (Glasauer & Shi 2022) to simulate possible neural representations of model variables. An obvious candidate is estimated elapsed stimulus duration, which is shown for different combinations of behavioral biases in Fig. 1C3 – 1E3. As possible correlate for expectancy we chose the prior probability density for stimulus duration (Fig. 1C4 – 1E4), which has a maximum close to 1.5 s, i.e., to the average stimulus interval. The temporal integral of this variable represents the instantaneous probability for stimulus offset: it increases from zero to one towards longer durations, because stimulus offset is expected almost with certainty for long enough stimulus durations. We used a weighted linear combination of these three variables, using log probabilities, to model the CNV (Fig. 7) by generating a simulated CNV response for each stimulus production phase and each participant, using the individual model parameters determined by fitting the model to the behavioral data of each participant (see also Fig. 3). The simulated CNVs were then averaged in the same way as the data CNV and plotted for comparison. While the frontocentral grand average CNV can be simulated quite well only by the expectancy (Fig. 6), when plotted separately for groups and previous stimuli, the match between CNV data and simulated CNV became correct only when using a weighted sum of all three variables (Fig. 7). A more positive simulated CNV is related to longer stimulus duration (Fig. 7A – 7C) and to higher central tendency (Fig. 7D – 7E), just as in the data. Correlation coefficients around 0.9 between predicted and experimental CNV time courses confirm the good match shown in Fig. 7 and helps to understand the main characteristics of the neural dynamics of the CNV and its dependence on previous stimulus and central tendency. For example, the CNV difference between low and high central tendency can be explained by the model, because for low CT (Fig. 1E) the internal estimate of elapsed time is almost linear with time (Fig. 1E3) and the prior expectancy is flat (Fig. 1E4), so that the sum of both will be smaller than the same sum for the case of high CT (Fig. 1D3 and 1D4).

Our Bayesian model also shows that prior information can directly affect the representation of elapsed time even when the stimulus is still ongoing: the weighting of the current temporal measurement with previous memory traces can happen instantaneously and does not have to wait until after the whole interval has been sensed and encoded. A similar conclusion has previously been drawn from behavioral evidence regarding the perception of visual tilt, which suggested that priors interact directly with early sensory signals (Cicchini et al., 2021).

We consider our current attempt of modeling the CNV a starting point for further model-based investigations. There are numerous possibilities that could be explored as possible correlates of model variables. For example, we assumed that log-probabilities would be represented in neural activity, as has been suggested previously (Haefner et al., 2024), but plain probabilities or other transformations would be possible as well. Furthermore, while our model represents a parsimonious solution to simulating individual combinations of central tendency and sequential dependence, more complex versions are possible, for example by including not just learning of the average stimulus duration, but also of the variability of the stimulus distribution (e.g., Piray and Daw, 2020).

There have been several attempts to explain the role of the CNV. The pacemaker-accumulator model of CNV, which links the CNV to absolute time accumulation (Macar et al., 1999; Macar and Vidal, 2004; Pfeuty et al., 2005; Casini and Vidal, 2011), has been questioned on the basis of previous findings supporting the view that CNV reflects temporal context (Wiener and Kanai, 2016; Li et al., 2017; Damsma et al., 2021; Baykan et al., 2024; Pang et al., 2024). Other theories posited the CNV as a marker of anticipation and preparation (Leuthold et al., 2004; Scheibe et al., 2009; Ng et al., 2011; Boehm et al., 2014). Here, we hypothesize that the CNV does not represent one single variable, but that it is a weighted sum of multiple signals related to estimation of elapsed time and expectancy. In summary, we propose our modeling as a method of how to relate the established function of the frontocentral cortex in time estimation and in anticipatory and preparatory processes (Pfeuty et al., 2005; Mento, 2013; Kononowicz et al., 2018) to quantifiable dynamics related to time perception and expectancy of stimulus offset.

In addition to the CNV during stimulus production we also investigated the two components of the postproduction stimulus-offset ERP, the P2 and N2 amplitudes. As shown previously (Kononowicz and van Rijn, 2014; Kruijne et al., 2021), both P2 and N2 showed a significant effect of current stimulus duration, suggesting a role in encoding temporal duration. N2, but not P2, was also related to sequential dependence at the group level. P2 showed a significant relation to central tendency at the group level and correlated, as mentioned above, with central tendency at an individual level. Notably, in terms of modeling, central tendency is inversely proportional to the steady-state weighting factors (Kalman gain) which determines how much weight is given to sensory information relative to prior information about the stimulus interval in the encoding phase. Thus, the relation between P2 and central tendency could reflect this weighting of prior information.

In conclusion, our study revealed neural correlates of individual perceptual biases reflected in the CNV and the postproduction offset ERP. The individual strategies of coping with uncertainty in time perception are thus represented already during the stimulus encoding phase, which suggests that perception of elapsed time is an ongoing process that does not require physically accurate temporal integration but that the integration process itself is directly affected by prior knowledge. As shown by the individual differences in sequential dependence, this prior knowledge incorporates previous stimuli in an inter-individually different manner, which is then reflected in the neural signatures of time estimation.

## Acknowledgments

The authors thank the cooperation of all participants in this study and Eduardo Martínez-Montes and Jhoanna Pérez Hidalgo-Gato for helpful discussions and comments on early versions of this work, and Juliane Pawlitzki for help with proofreading. Funding: Deutsche Forschungsgemeinschaft DFG grant Gl 342/3-2.

## Supplemental Material

### 1. Deriving an upper bound for sensory noise variance

The free parameters of the two-state Kalman filter model with the steady state equations given in Eqn. 2 are the variance ratios 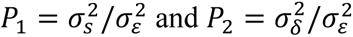. However, for the prior probability density of stimulus duration 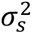 knowledge of the variance 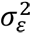 is required. We therefore must estimate the unknown noise variance 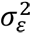. Here we derive this variance for the two boundary cases of the two-state model in the linear domain. To do so, we assume that the observed residual variance after fitting the model is due to the unknown sensory noise. With stimuli *s*, model outputs *ŝ*, and data *d*, the observed reproduced values, the model residual is *r* = *ŝ* − *d*, the residual variance is 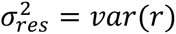.

For the simple static model, *K*_2_ = 0. We can set the mean of the stimulus distribution to zero, *E*[*s*] = *m*_*i*_ = *m* = 0 without loss of generality. The static model estimate is thus *ŝ*_*i*_ = *K*_1_*s*_*i*_. From the data values we get *d*_*i*_ = *K*_1_*x*_*i*_ = *K*_1_(*s*_*i*_ + *ε*). Thus, the residual is *r* = *ŝ* − *d* = *K*_1_*s* − *K*_1_(*s* + *ε*_*s*_) = −*K*_1_*ε*_*s*_, and therefore 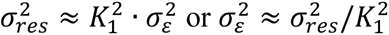. This also corresponds to numerical simulations.

For *K*_2_ = *K*_1_ the model reduces to the simple iterative model *ŝ*_*i*_ = *K*_1_*s*_*i*_ + (1 − *K*_1_)*ŝ*_*i*−1_. The corresponding equation for the data is *d*_*i*_ = *K*_1_(*s*_*i*_ + *ε*) + (1 − *K*_1_)*d*_*i*−1_. The residuals are

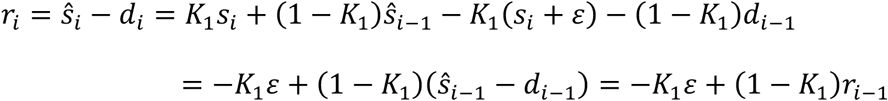

Thus *r*_*i*_ − (1 − *K*_1_)*r*_*i*−1_ = −*K*_1_*ε*. Therefore we can approximate the variance 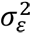 as *var*((*r_i_* − (1 − *K*_1_)*r*_*i*−1_)/*K*_1_), from which follows 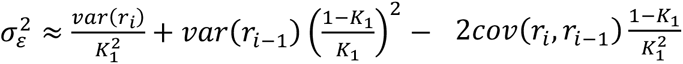, because the residuals are autocorrelated due to the noise propagating through the time series. The covariance of the residuals due to the noise propagation is *cov*(*r*_*i*_, *r*_*i*−1_) = (1 − *K*_1_)*var*(*r*_*i*_). Therefore, we can approximate 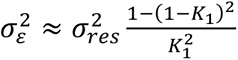.

In both cases, this is an upper bound, because the observed variance is likely a combination of execution noise and sensory noise, rather than being only due to sensory noise. With the estimated variance 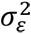 we can now calculate from the fitted parameter *P*_1_ the variance of the prior distribution of stimulus duration as 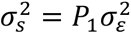.

The instantaneous expectancy for stimulus offset at trial *i* over time, expressed as prior probability density of stimulus duration, is accordingly

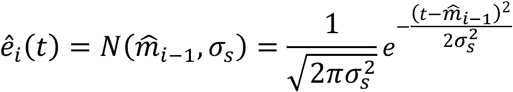

The temporal integral 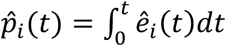 is thus the cumulative prior density and represents the instantaneous probability for stimulus offset over time at trial *i*. It is zero at stimulus onset and increases towards 1 with stimulus duration.

### 2. Left parietal CNV responses

**Figure S1.**
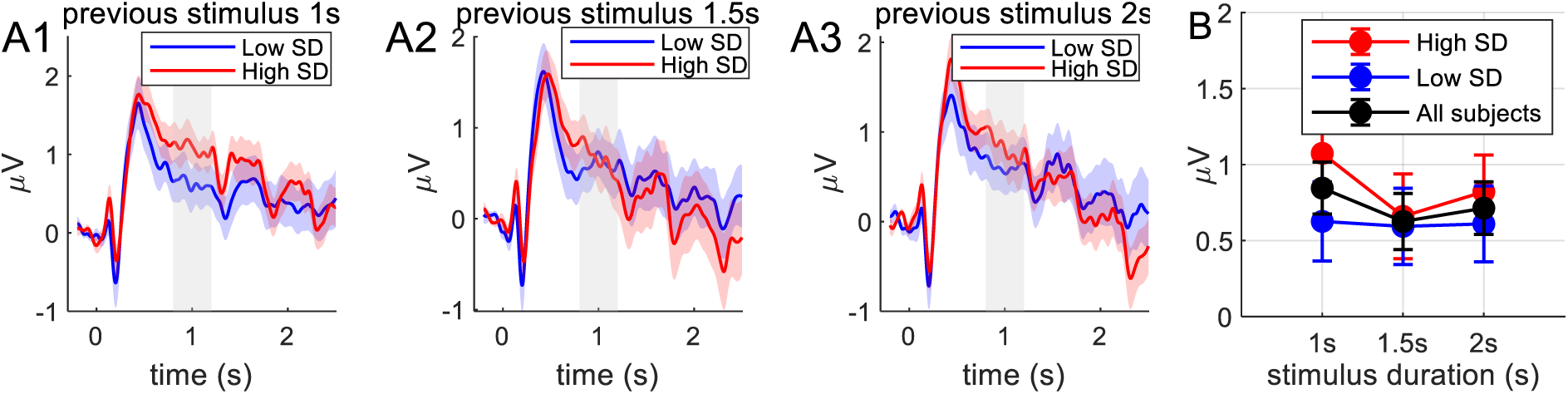
Left parietal grand average CNV responses. A1-A3: CNV response grouped by previous stimulus separately for previous stimulus (A1:1s, A2:1.5s and A3:2s) during the production phase recorded from the left parietal electrodes (P3, P7), comparing subjects grouped by sequential dependence with High SD (blue, n=23) and low SD (red, n=23). B. average CNV amplitudes in the time interval of 0.8–1.2 s (gray shaded region in time-course plots) for low SD (blue), high SD (red), and all subjects (black). Error bars represent the standard error of the mean. Unlike the right parietal region, no systematic differences between high and low SD groups were observed in the left parietal region (F(1,43)=0.40, p=0.40, η^2^=0.01) supporting the specificity of sequential dependence effects to the right hemisphere, and also CNV amplitude did not differ significantly over the stimulus duration (F(2,86)=0.8, p=0.45, η^2^=0.006) with no interaction with the sequential dependencies groups (F(2,86)=0.58, p=0.55, η^2^=0.004). Furthermore, repeated measures ANOVA comparing CNV amplitudes between left and right parietal regions with main factor of region and between-subject factor of Sequential Dependence (SD) revealed no significant main effect of region (F(1,43)=0.679, p=0.414, η²=0.004). However, a significant interaction between region and SD was observed (F(1,43)=5.437, p=0.024, η²=0.028), indicating that the difference in CNV amplitudes between parietal regions depended on sequential dependence grouping. Additionally, there was a significant main effect of SD group (F(1,43)=5.008, p=0.030, η²=0.078). Bonferroni-corrected post hoc comparisons showed that the High SD group exhibited significantly greater CNV amplitude only in the right parietal region compared to the Low SD group (t=3.105, p=0.017). No other significant differences were observed in post hoc comparisons between groups or regions (all other p-values >0.10).

### 3. Parietal production offset responses

**Figure S2.**
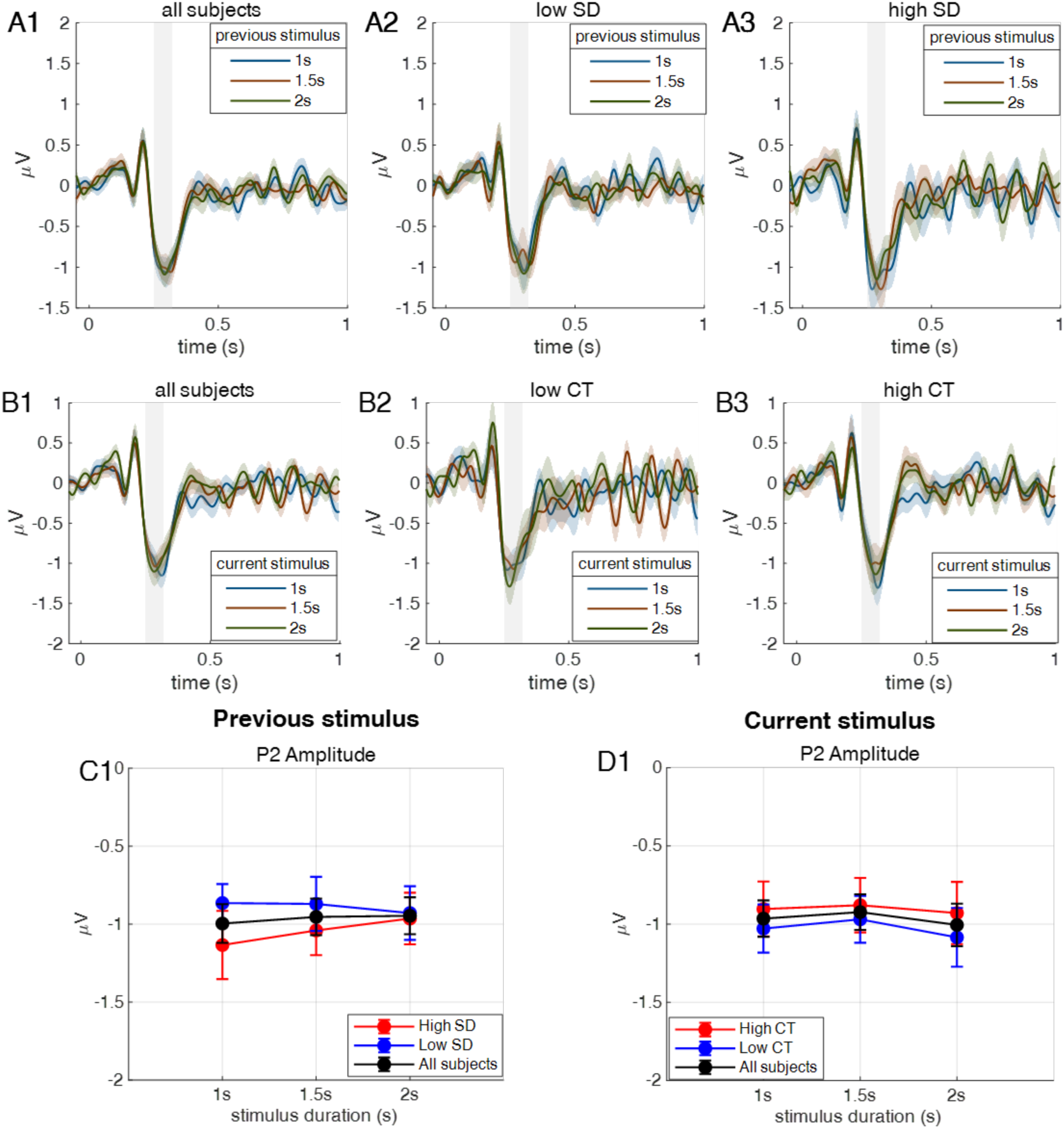
Right Parietal postproduction offset ERP responses. **A1:** Grand-average ERP waveforms from the right parietal electrodes, separated according to the duration of the previous trial stimulus for all subjects (n=45). **A2 and A3:** ERP responses similarly grouped by previous trial stimulus durations, plotted separately for participants with low sequential dependence (Low SD, n=23; A2) and high sequential dependence (High SD, n=22; A3). **B1–B3:** ERP waveforms plotted by current trial stimulus duration for all subjects (B1), low central tendency (Low CT, n=23; B2), and high central tendency (High CT, n=22; B3) groups. **C1:** Postproduction P2 amplitude, computed as the mean within the time interval indicated by the gray-shaded region, plotted against previous stimulus duration for Low SD (blue), High SD (red), and all subjects (black). A repeated measures ANOVA indicated no significant effects of previous stimulus duration (F(2,86)=0.21, p=0.81, η²=0.0008) or sequential dependence grouping (F(2,43)=0.52, p=0.47, η²=0.01), and no significant interaction between these factors (F(2,86)=0.85, p=0.42, η²=0.003). **C2:** Mean P2 amplitude plotted against the current stimulus duration for participants with low CT (blue), high CT (red), and all participants (black). Similarly, repeated measures ANOVA showed no significant effect of current stimulus duration (F(2,86)=0.39, p=0.67, η²=0.002), no differences between CT groups (F(1,43)=0.68, p=0.41, η²=0.013), and no interaction effect between stimulus duration and CT group (F(2,86)=1.15, p=0.3, η²=0.005).

**Figure S3.**
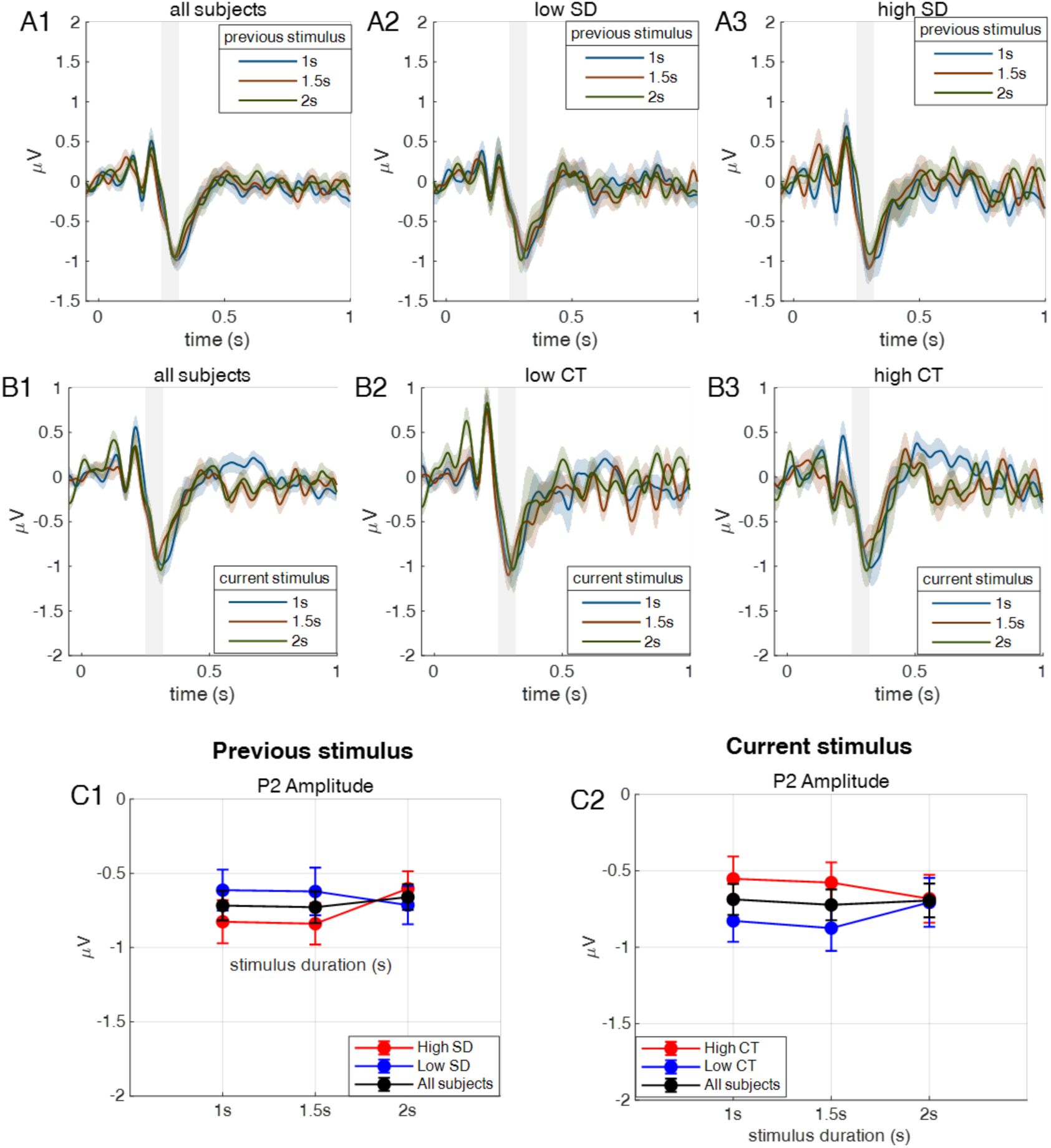
Left Parietal postproduction offset ERP. **A1**: Grand average response of postproduction offset ERP for all subjects (n=45) grouped based on previous trial stimulus. **A2**. For low SD group (n=23). **A3**. For high SD group (n=22). **B1-B3**. Postproduction ERP plotted grouped based on current trial stimulus duration (**B1**: all subjects, **B2:** low CT, **B3**: high CT). **C1.** Postproduction P2 amplitude plotted over previous stimulus duration for subjects with low SD (blue), high SD (red), and all subjects (black). A repeated measures ANOVA showed no main effects for main factors of *previous stimulus* duration (F(2,86)=0.21, p=0.37, η^2^=0.002) and sequential dependence *group* (F(1,43)=0.40, p=0.47, η^2^=0.007) on CNV amplitude, calculated as the mean value within P2 amplitude time interval indicated by gray region. Also, there was no interaction(F(2,86)=2.204, p=0.11, η^2^=0.014). **C2:** N2 amplitude plotted over current stimulus duration for subjects with low CT (blue), high CT (red), and all subjects (black). Repeated measures ANOVA, revealed no differences for current stimulus duration (F(2,86)=0.051, p=0.95, η^2^=0.0004) nor between the central tendency groups duration (F(1,43)=0.39, p=0.53, η^2^=0.006) with no interaction between the current stimulus and central tendencies groups (F(2,86)=0.48, p=0.3, η^2^=0.004).

